# Parsing out the variability of transmission at central synapses using optical quantal analysis

**DOI:** 10.1101/624692

**Authors:** Cary Soares, Daniel Trotter, André Longtin, Jean-Claude Béïque, Richard Naud

## Abstract

Properties of synaptic release dictates the core of information transfer in neural circuits. Despite decades of technical and theoretical advances, distinguishing bona fide information content from the multiple sources of synaptic variability remains a challenging problem. Here, we employed a combination of computational approaches with cellular electrophysiology, two-photon uncaging of MNI-Glutamate and imaging at single synapses. We describe and calibrate the use of the fluorescent glutamate sensor iGluSnFR and found that its kinetic profile is close to that of AMPA receptors, therefore providing several distinct advantages over slower methods relying on NMDA receptor activation (i.e., chemical or genetically encoded Calcium indicators). Using an array of statistical methods, we further developed, and validated on surrogate data, an expectation-maximization algorithm that, by biophysically constraining release variability, extracts the quantal parameters n (maximum number of released vesicles) and p (unitary probability of release) from single-synapse iGluSnFR-mediated transients. Together, we present a generalizable mathematical formalism which, when applied to optical recordings, paves the way to an increasingly precise investigation of information transfer at central synapses.

## I. INTRODUCTION

Our understanding of the factors that contribute to the stochastic and variable process of synaptic transmission has improved steadily over the last few decades [10, 28, 40]. It is now generally agreed that, at most glutamatergic synapses, quantal release does not saturate postsynaptic receptors [26, 27, 32, 33, 49] and that variability in trial-to-trial neurotransmission arises primarily from differences in the profile of glutamate released into the synaptic cleft [40]. Several presynaptic mechanisms have been proposed to account for such amplitude fluctuations – uneven packaging of glutamate into synaptic vesicles, differences in release location within a synaptic terminal, diffusion process in the synaptic cleft and mode of exocytosis [12, 19, 41, 52]. As an additional factor, multiquantal release has been observed at many central synapses [1, 13, 15, 22, 35, 42, 50], where two or more vesicles are released quasi simultaneously at single synapses in response to the same electrical stimulus. Since each of these sources of variability impact the transmission of information differently, it is therefore important to parse out the relative proportion of different sources of variability at central synapses.

Several experimental methodologies have been developed to monitor transmission at single synapses [17, 18, 30, 35, 42]. Here we describe an optical-based technique and provide a number of validation and calibration experiments for the intensity-based optical glutamate sensor, iGluSnFR [31], for optical quantal analysis at central synapses. We further provide a detailed theoretical and quantitative analysis for estimating fundamental features of quantal glutamate release. Leveraging experimental and statistical techniques, combined with a theoretically sound model, we present a formalism that is well poised to parse out the structure of variability and information content at central synapses.

## II. RESULTS

To study quantal features of glutamate release at central synapses, we turned to a genetically encoded intensity-based glutamate sensing florescent reporter (iGluSnFR). The versatility and usefulness of iGluSnFR as an optical reporter of glutamate release has been demonstrated in both microscopic and macroscopic brain compartments [9, 31, 36, 37, 53], although it has relatively seldom been used to study features of glutamate release at single spines [45]. To this end, we introduced iGluSnFR along with the morphological marker mCherry to CA1 neurons in hippocampal organotypic slices using biolistic transfection several days prior to the experiments (Figure 1A) [45, 46]. A detailed description of these procedures is available in [46]. This overall approach was favoured since it allows for sparse transfection thereby allowing us to resolve optical signals from single spines with high contrast. Transfected neurons were imaged by two-photon microscopy using an excitation wavelength of 950 nm (Figure 1B) which we found to allow detection of both the iGluSnFR and mCherry fluorescent signal simultaneously. Dendritic spines in the apical dendritic arbor of transfected CA1 neurons were targeted for optical quantal analysis experiments. These contacts are likely the postsynaptic targets of Schaffer’s collateral axons.

**FIG. 1.**
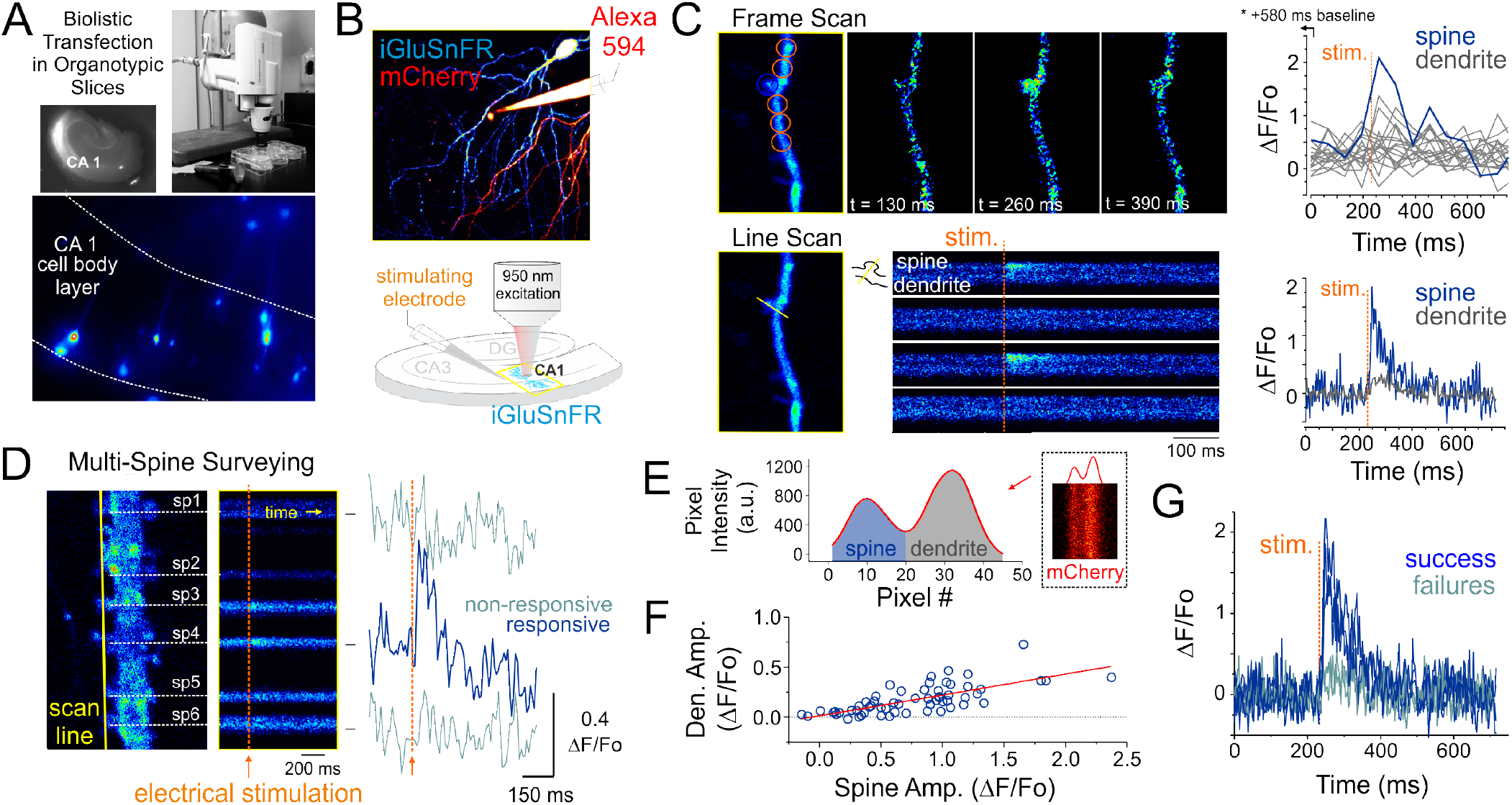
Optical detection of glutamate release at single synapse using an genetically encoded glutamate sensor **A** Biolistic transfection of CA1 hippocampal neurons with the intensity based glutamate sensor, iGluSnFR. **B** Experimental setup. A glass electrode filled with a fluorescent dye (Alexa 594) was positioned in the stratum radiatum adjacent to an iGluSnFR-expressing cell and was used to deliver electrical stimuli to the slice to evoke endogenous glutamate release. Neurons were typically transfected with both iGluSnFR and mCherry and expressed variable amount of the fluorescent proteins. **C** A comparison of iGluSNFR transients recorded at the same spine using either a frame-scan (spatial resolution > time resolution) or a line-scan (time resolution > spatial resolution) configuration. The fastest frame scan sampling rate of our optical system is 65 ms per frame, whereas rates of ≈ 1.4 ms perline were typically obtained in line-scan mode. **D** A line scan experiment is shown where multiple adjacent spines are surveyed simultaneously for evoked iGluSnFR transients. Shown at right is the time series resulting from a continuous line scan before and after an electrical stimulation. Spines 3 and 4 showed responsiveness to the electrical stimulus in this trial. **E** Isolation of spine and dendrite specific signals from a line scan is achieved by averaging pixels in their respective compartments, which was inferred by the presence of a dip in the mCherry signal. **F** The amplitude of spine iGluSnFR signals (same spine as in panel D) plotted against the corresponding amplitude of dendritic signals. A linear regression results in a significant positive correlation (slope 0.21, adjusted R-squared = 0.496). G Clearly distinguishable spine successes and failures demonstrate the probabilistic nature of vesicular release at these synapses.

### A. iGluSnFR-mediated monitoring of endogenous glutamate release

Pyramidal neurons were identified by their localization in the slice and morphology. Namely imaging targeted to the CA1 region and we sought the clear presence of basal and apical spinous dendritic arborisation. The morphological identification was typically carried out by solely monitoring mCherry fluorescence. However, the baseline iGluSnFR fluorescence was typically fairly high, homogenously distributed across neuronal compartments and spines were readily observable, thereby readily allowing for broad cell-type identification. A typical experiment began by randomly surveying the apical arbor of an iGluSnFR-expressing cell for dendritic spines that exhibit a time-locked fluorescent responses to electrical stimuli delivered via a glass pipette positioned in stratum radiatum. In a few experiments, Alexa 594 was included in the internal solution of the stimulating electrode for direct visualization (Figure 1B), however, in the majority of experiments this dye was omitted and the stimulating electrode was maneuvered in the slice under visual guidance solely using differential interference contrast microscopy.

The optical detection of synaptic events that are eminently short-lived, spatially distributed and scarce is inherently challenging and deserves attention. In principle, imaging in frame scanning mode would be ideal to monitor synaptic fluorescent events from large dendritic regions, but it is hindered by limited signal to noise ratio and temporal resolution (Figure 1C). We thus carried out line scan experiments wherein multiple neighboring spines were monitored simultaneously (Figure 1D). This approach offered the ability to survey multiple spines at once with a scanning frequency (>500 Hz) sufficient to visually identify rapid glutamate transients. To circumvent the relative paucity of synaptic events due to the probabilistic nature of release, paired-pulse electrical stimulation (50-100 ms inter-stimulus interval) were delivered to increase the likelihood of release. Lastly, a realistic range of stimulus intensities was determined by parallel and historical whole-cell electrophysiological recordings by the same experimenter. Once a responsive spine was identified, a short line scan was redrawn through the spine and its parent dendritic compartment to capture the spatial profile of glutamate release. The electrical stimulation was then gradually reduced to the minimal intensity that still evoked time-locked responsiveness. This last step was taken in order to reduce the potential of signal contamination by glutamate spillover from neighbouring synapses. The identified spines routinely stayed responsive to electrical stimulation for long durations (> 1 hour), opening the door to the repetitive low frequency sampling methodology required for building a dataset sufficient for optical quantal analysis.

### B. Extraction of regions of interest

Spatial discrimination of iGluSnFR signals emanating from either spine or dendritic compartments was achieved by analyzing the intensity profile across the line scan, which was drawn orthogonal to the parent dendrite. The trough between spine and dendrite peaks was used to split the signal of the line scan into the two compartments (Figure 1E) to isolate spine- and dendrite-specific iGluSnFR transients (Figure 1D, right). Larger amplitude iGluSnFR transients were generally observed in the spine compartment, indicating that the density of glutamate release was mostly concentrated at the spine. When present, the dendritic fluorescence transients were of smaller amplitudes and co-varied with that recorded from the spine compartment, suggesting that dendritic signals were likely the result of spillover from the parent spine rather than from release from a distinct, neighbouring synapse (Figure 1F). As such, we used only the spine compartment signal for all subsequent analyses. Finally, and consistent with the probabilistic release of glutamate vesicles at these synapses, release failures were readily observed (Figure 1G). These results demonstrate that iGluSnFR is a useful optical reporter for single-spine quantal analysis.

### C. Glutamate vs post-synaptic calcium sensors for opto-quantal analysis

A difficulty in unambiguously and routinely study neurotransmitter release from a single synapse arises from the lack of spatial resolution afforded by electrophysiological recordings. By providing spatial information, optically-based approaches for quantal analysis offers promise of a solution to this problem, yet are limited by temporal resolution generally poorer than that afforded by cellular electrophysiology. By using two-photon uncaging of MNI-glutamate to precisely control the amount and timing of glutamate released onto single spines, we next sought to examine the kinetic performances of iGluSnFR by a side-by-side comparison with other commonly used reporters of glutamate release for quantal analysis. Specifically, we sought to compare electrophysiological monitoring of synaptic AMPAR activation and optical recordings of quantal analysis using NMDAR-mediated calcium influx by calcium indicators. Since optical recordings of calcium influx using the GCaMP family of genetically encoded calcium-indicators are becoming increasingly popular, we turned our attention to GCaMP6f, a fast variant of the GCaMP family.

We obtained whole-cell recordings from CA1 neurons transfected with either iGluSnFR of GCaMP6f (Figure IICA) and voltage-clamped the cell at −70 mV. While continuously imaging the spine of interest (at ≈ 715 Hz), a second laser line tuned to 720 nm delivered a 1 ms light pulse to the tip of the spine in the presence of MNI-Glutamate (2.5mM), to induce uncaging-evoked optical transients recorded simultaneously with EPSCs (Figure IICB-C). In response to repetitive presentation of nominally constant concentration of glutamate by 2P uncaging at single synapses (Figure IICD), we compared the performance of 3 distinct reporters of glutamate transients at single synapses: i) iGluSnFR transients; ii) GCaMP6f transients (i.e., NMDAR-dependent calcium influx) and, iii) AMPAR activation (uncaging-evoked EPSCs; uEPSCs). We found that the trial-to-trial variability of the iGluSnFR responses was remarkably low, even lower than that of uEPSCs (Figure IICE). In keeping with the more complex and convolved nature of the NMDAR- and calcium-mediated optical detection of glutamate release, the GCaMP6f signal displayed the largest variability of the 3 approaches (Figure II CE). iGluSnFR transients also displayed much faster decay kinetics (Figure IICF) and rise time (Figure II CG) compared to GCaMP6f (p<0.001, unpaired student’s t-test), and were remarkably close to the kinetics of uEPSCs. The kinetic properties of iGluSnFR in response to glutamate uncaging therefore favorably compares to those of the calcium-sensitive organic dyes Alexa 4FF (Lee et al., 2016) and Oregon Green BAPTA-1 (unpublished observations) that are significantly slower. The fast kinetics of iGluSnFR enable discrimination of successive stimulus peaks at higher stimulus frequencies (50-100 ms Interstimulus interval; Figure II CH) without the need of signal deconvolution. Moreover, neither the amplitude nor the kinetics of the iGluSnFR responses were affected by the competitive AMPA receptor antagonist NBQX (*n* = 4, Figure II CI), indicating that iGluSnFR-based quantal analysis could be performed in the presence of a glutamate receptor antagonists. This possibility may offer some flexibility to avoid specific experimental complications, such as minimizing excitability for experiments in highly recurrent networks or minimize plasticity induction by repetitive and prolonged stimulation paradigms. Altogether, the iGluSnFR method for quantal analysis offers more experimental flexibility and faster kinetics than that afforded by NMDAR-mediated Calcium influx detected by GCaMP6f, with a temporal resolution that nears that of cellular electrophysiology.

### D. Biophysics of glutamate release variability

The goal of quantal analysis is to infer release properties of glutamate release from a distribution of recorded release magnitudes. Quantal analysis of synaptic release has been performed for decades and the formalism has evolved and adapted as new and improved recording technologies were developed. For didactic purposes, we revisited here some of the basic assumptions commonly held for performing quantal analysis of glutamate release events at single synapses. We started exploring the most appropriate continuous distribution to describe the inherent variability expected of a glutamate quantum. Our aim was to derive from the biophysical features of synaptic vesicles a mathematical description of the expected distribution of glutamate release amplitudes, along with their expected variability.

Based on previous theoretical studies, we expect that the variability of inner vesicular volumes (Figure 3A) will be a potent determinant of the variability in the amount of glutamate molecules per quantum [4]. What variability of glutamate release do we expect from fluctuations in vesicle diameters only? In order to find this, we first constrain the concentration of glutamate within synaptic vesicles to be constant across the many synaptic vesicles of a given neuron. Next, we assume faithful release of a single vesicle and that the relative location and loading of vesicles does not introduce a significant amount of variability in the activation of post-synaptic receptors. We will revisit these assumptions sequentially as we assemble the mathematical synapse model.

**FIG. 2.**
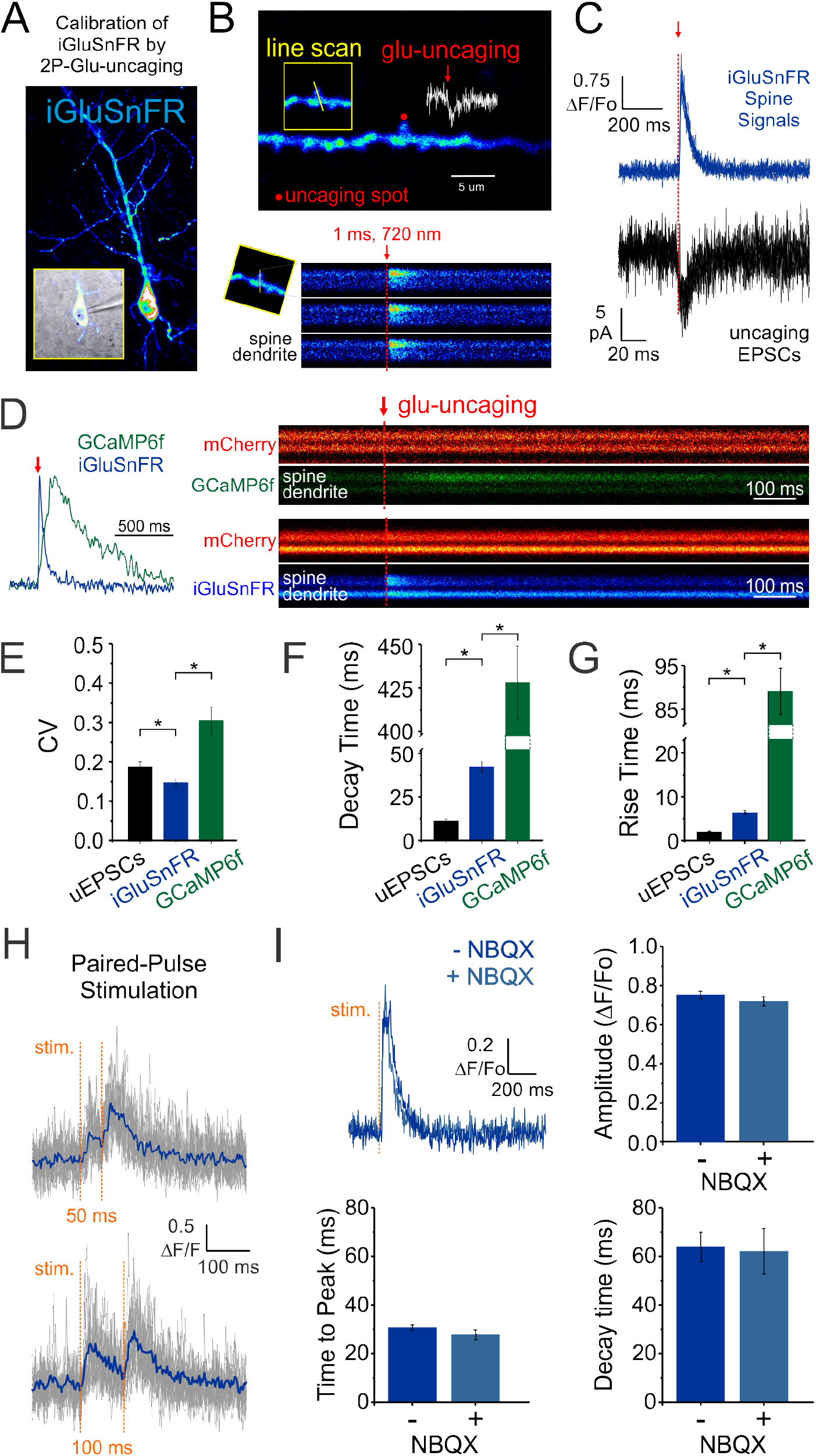
Features of iGluSnFR-mediated responses. **A** Whole-cell recording of an IGluSnFR-expressing CA1 neuron. **B** Optically-evoked iGluSnFR transients were generated at a single spines by two-photon uncaging of MNI-glutamate. A continuous line scan was imaged at 950 nm while a second laser line tuned at 720 nm was used to deliver the uncaging events (1 ms; red arrow). **C** Spine iGluSnFR fluorescence transients from 10 consecutive stimuli are displayed on the right panel, along with the corresponding uncaging evoked EPSCs. **D** A kinetic comparison of iGluSnFR and GCaMP6. In response to an identical 1 ms light pulse used to uncage MNI-glutamate at a single spines, the decay (**F**) and rise (**G**) kinetics of iGluSNFR transients were much faster than calcium transients recorded from GCaMP6f-transfected neurons, but still slower than corresponding uncaging-evoked excitatory postsynaptic currents. **H** The rapid kinetics of iGluSNFR enables peak-detection at stimulation frequencies that are suitable for studying synaptic facilitation and depression. **I** NBQX, an antagonist of AMPA-type glutamate receptors, has no effect on the amplitude or kinetics of evoked iGluSnFR transients (n = 50 stimuli in each condition; p > 0.05 in all cases, paired students t-test) and can be administered during optical quantal analysis to dampen plasticity that might be induced by repetitive sampling. Time to peak (after stimulus) was used to quantify rise times in this scenario rather than the 20-80% rise time method used previously on uncaging-evoked iGluSNFR transients (**G**) since a subset of evoked transients with small amplitudes were significantly impacted by optical noise leading to misleading measurements using the 2080% rise time method.

**FIG. 3.**
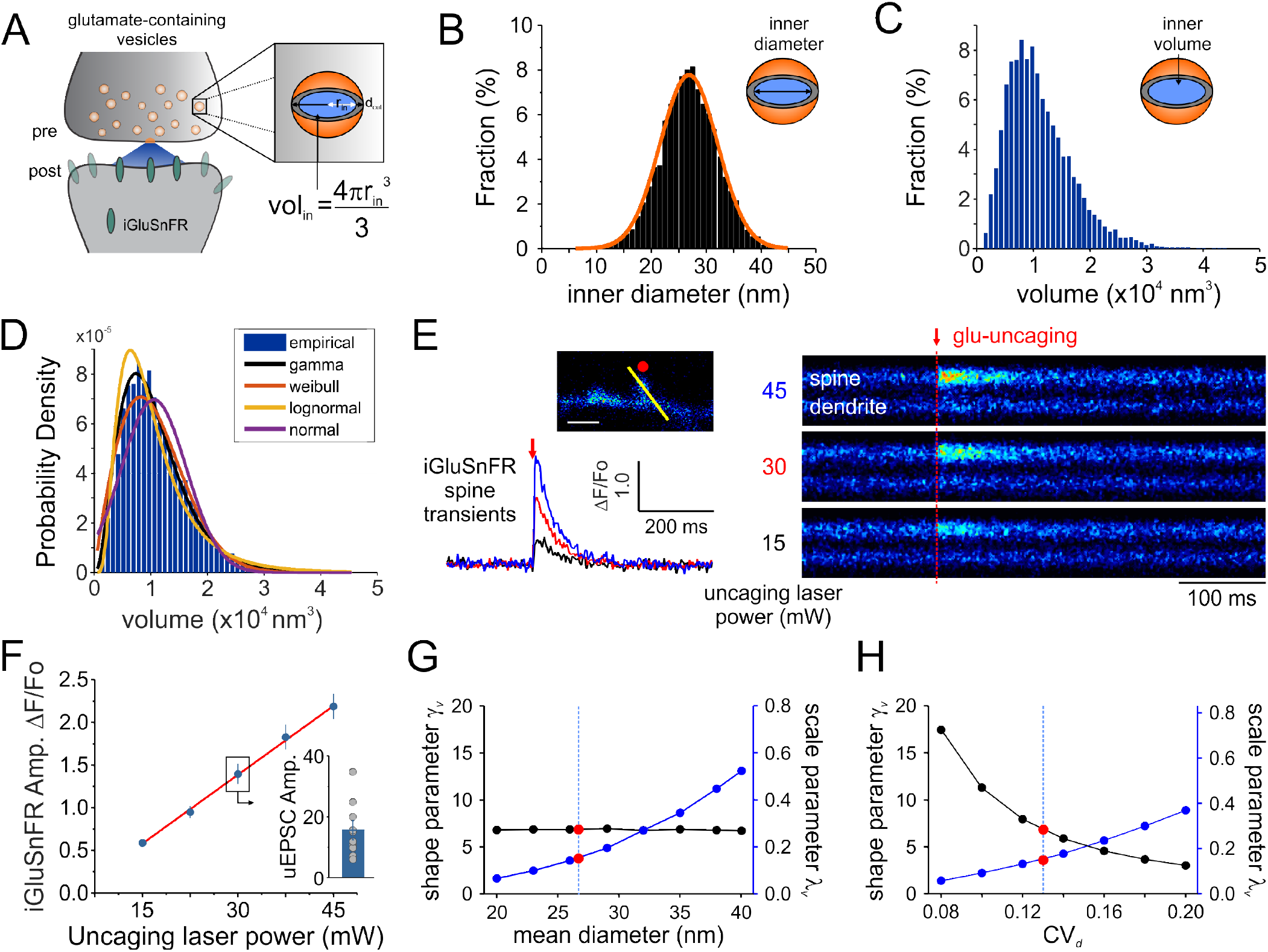
Modelling Synaptic Glutamate Transients Following Vesicle Release. **A** A schematic description of the measurements and calculation used to infer synaptic vesicle volumes. This analysis was based on the assumptions that: i) the distribution of synaptic vesicle diameters is uniform and; ii) the shape of a synaptic vesicle is roughly spherical. **B** A theoretical distribution of synaptic vesicle diameters using electron microscopy measurements described in Qu et al. 2009 (outer vesicle diameter = 38.7 nm; CV = 0.13; n = 10,000 vesicles, [38]). The inner diameter of SVs was calculated by subtracting the thickness of the vesicular membrane (2×6 nm). **C** A distribution of the inner volume of SVs assuming that each is approximated by the volume of a sphere. **D** Fitting of various continuous distributions to the modeled volume distribution, ordered in the legend based on the Bayesian Information Criteria (BIC). **E** iGluSnFR transients generated at a single spine by two-photon glutamate uncaging at different uncaging laser powers (mW = milliwatts of power after the objective). **E** A positive linear relationship between uncaging laser power and the amplitude of iGluSNFR transients at a single spines (Adjusted *r*^2^ = 0.997) indicates that the transduction is linear within this range. **G-H** Thoertical relationship between the parameters of a gamma distribution and the properties of the vesicle dimensions. Shape and scale parameter values are shown against the mean vesicle diameter for CVv = 0.13 (**G**) and as a function of CVv for mean vesicle diameter of 38.7 nm (**H**).

Electron microscopy studies have shown that the variability in vesicle diameters at hippocampal synapses is normally distributed. Using the measurements obtained from one such study (mean vesicle diameter 38.7 nm, CV*_d_* = 0.13 [38]) we generated a simulated distribution of 10,000 inner vesicle diameters (Figure 3B) and a corresponding distribution of the inner vesicle volumes (Figure 3C), assuming the shape of synaptic vesicles is approximated by a sphere. Inner vesicle diameters and volumes were calculated by first subtracting the thickness of the plasma membrane (12 nm [38]). This volume distribution can be readily calculated by a change in variable of the diameter distribution [2, 4]. In line with the cubic relationship between volume and diameter, the resulting distribution (Fig. 3C) is non-Gaussian as it displays an important rightward skew.

To compare the possible distributions of volumes emanating from a range of experimentally derived vesicular diameter, we explored a set of continuous distributions (normal, gamma, Weibull, lognormal) that could accurately describe the skewed distribution of inner vesicle volumes simulated. Using the Bayesian Information Criteria (BIC) as a scoring metric, we could rank the distributions with their degree to which they fit the simulated distribution (Figure 3D). We found that the gamma distribution provided the best approximation of the empirical distribution of vesicle volumes, *p*(*v*), followed by the Weibull, lognormal, and finally the normal distribution. This finding is intriguing when we consider that many previous studies of quantal analysis have reported using a Gaussian mixture model of release events [21, 23, 29], although at least one study has used skewed distributions [2] and at least one study a gamma distribution [6].

The gamma distribution is described by two parameters: a shape parameter *γ* and a scale parameter λ and it is expressed in terms of the gamma function Γ(·). When used to approximate the distribution of vesicle volumes arising from normally distributed diameters, we write

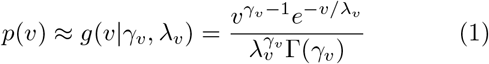

where λ*_v_* and *γ_v_* are the parameters for the volume distribution. These paremeter values can be found by matching the first two moments of simulated (Fig. 3) and theoretical (Eq. 1) distributions. Equation 1 has a right-skew controlled by the parameter *γ_v_*. Conveniently, its mean (*E*[*v*] = *γ_v_*λ*_v_*), its variance 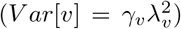, its skewness 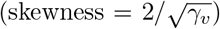 and its coefficient of variation 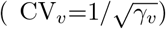 are simple expressions of these parameters. Also of considerable practicality, the addition of two independent gamma-distributed variables results in a random variable that is itself gamma-distributed with shape parameter equal to the sum of the shape parameters. As pointed out by Bhumbra and Beato (2013) [6], these properties allow for a clearer treatment glutamate release variability without explicitly compromising the validity of the gamma-release model.

To relate the parameters of the gamma distribution with vesicle dimensions, we calculated the expected range of *γ_v_* and λ*_v_* as a function of the mean vesicle diameter (*μ_d_*, Figure 3E) and diameter coefficient of variation (CV*_d_*; Figure 3F) for simulated vesicle volume distributions. The shape parameter is unaffected by changes in mean diameter, but the scale parameter increases nonlinearly with increasing diameters. In addition, the shape parameter decreases and the scale parameter increases when the CV of vesicle diameter increases. It is therefore possible to interpret an increase of the scale parameter as an increase in the mean vesicle diameter, but only if the shape parameter shows no concomitant changes.

What are the theoretical predictions of vesicle volume variability for optical measurements of cleft glutamate? Using the mean diameter *μ_d_* and the variability of diameters CV*_d_* from electron microscopy recordings, we predict λ*_v_* = 0.15 and *γ_v_* = 6.8. Importantly, these parameters give rise to a variability of volumes CV*_v_* of 0.38. In theory, unequal loading, diffusion and observational noise should increase the coefficient of variation once we consider the glutamate reported on the post-synaptic membrane instead of vesicle volumes. Since these factors are likely to be captured by another skewed distribution [6, 8, 19] such as the gamma distribution, it is appropriate to use a gamma distribution to capture the total variability of univesicular releases. To consider a possible discrepancy between the variability of univesicular releases and that of vesicle volumes, we use *γ* and λ to parameterize the distribution of univesicular releases, not to be confused with the parameters of the theoretical volume distribution *γ_v_* and λ*_v_*. In fact, since additional source of variability can only increase the CV, our *γ_v_* should be considered an upper bound on *γ*. To summarize, we derived biophysical constraints for the parameters of a gamma-distributed set of univesicular glutamate release events (UVR) using previous measurements of the distribution of inner vesicular volumes and the assumption of equal glutamate concentration across vesicles.

### E. Observational error and iGluSNFR transduction

In principle, the experimental readout expected from the non-uniform distribution of cleft glutamate will arise in part from the cubic transform outlined above but it can be corrupted by loading, diffusion and by non-optimality of the iGluSnFR signal transform. In order to begin addressing the issue of iGluSnFR transform, we sought to experimentally interrogate as directly as possible the relationship between the quantity of glutamate release at single spines and the amplitude of iGluSNFR-mediated transients. By varying the amount of glutamate released onto dendritic spines through step-wise increments in uncaging laser power during simultaneous optical and electrophysiological recordings (Figure 3E), we found that the relationship between uncaging laser power and iGluSnFR amplitude was linear (Figure 3F) within the expected physiological range of glutamate release, as determined by the average amplitude of uEPSCs [3, 25, 45, 46]. Thus, the iGluSNFR-mediated optical signal appears to linearly report glutamate concentration.

We then estimated a convolved metric of observational error CV*_opt_* to be 0.15, by measuring the variability of the iGluSNFR transients upon presentation of nominally fixed amounts of glutamate concentrations following repetitive uncaging at a fixed laser intensity (around 30 mW; Figure IICE); while uEPSC amplitudes were within an expected physiological range (Figure 3E). At most, adding this measurement noise brings the combination of diameter and optical variability to 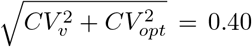. The formalism outlined above therefore predicts the distribution of optical signals when glutamate is released from a presynaptic terminal. We next considered the variability imposed by the stochastic nature of vesicle releases.

### F. Release failures

A large part of the variability is attributed to the stochastic failure of vesicle release [11] upon action potential arrival. To formally include this process in our predicted distribution of optical signals, we considered a mixture model wherein we stochastically sampled from one of many independent sub-distributions, which are called components. Since in certain conditions, multiple vesicle release (MVR) occurs at central synapses [13, 15, 35, 50], we consider a MVR model for which uni-vesicular release (UVR) is a special case. When *n* vesicles are docked and ready to be released and when each of these vesicles is released independently with probability *p*, the number of vesicles released will follow the binomial distribution. At times, we will expect that all vesicles have failed to release, in which case we will sample from the failure distribution. Assuming a Gaussian measurement (here called optical) noise for the failure distribution, we obtain the gamma-Gaussian mixture

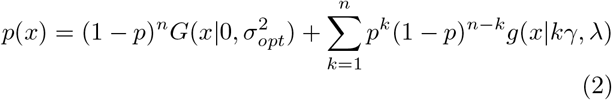

where 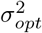 is the variance of the optical noise, *G*(*x*|*μ*, *σ*^2^) is a Gaussian distribution and *x* is an observation of cleft glutamate here in units of fluorescence Δ*F*/*F*_0_ but could be generalized to measures of post-synaptic current amplitudes. In Eq. 2, *k* ranges from 1 to *n* and refers to the possible number of vesicle released. The binomial coefficient *p^k^*(1 − *p*)^*n−k*^ establishes the probability of observing *k* vesicles, while each time that *k* vesicles are released, the cleft glutamate is obtained as a sample from a gamma distribution *g*(*x*|*kγ*, λ) with a shape parameter corresponding to *k* times the univesicular shape parameter *γ*.

We make three observation on this gamma-Gaussian mixture. Firstly, we distinguish the vesicular release probability p from the probability of any vesicle being released *P* = 1 − (1 − *p*)*^n^*. Secondly, the mean and the variance of this distribution now depend on the maximum number of vesicles released *n*, namely *μ* = *npγ*λ and 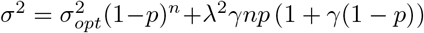. Lastly, it can be useful to analyze the measured variability, *CV*, in terms of the variability of univesicular releases 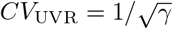, the variability due to observational error *σ_opt_* and the variability of a binomial process 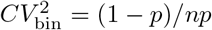. In this way we can parse the variability in three terms

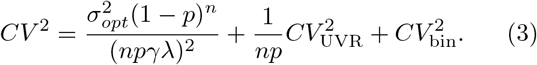

This expression allows us to parse out the variability in terms of distinct sources.

Overall, for the experiments described in Fig. 1, the gamma-Gaussian mixture should capture the variability of glutamate-dependent optical events originating from:

1. Optical: various optical measurement noise,
2. Binomial: the stochastic behaviour of releasing *n* vesicle independently,
3. Unitary: release variability associated with each vesicle release.

The latter comprises variability from vesicle sizes, loading and diffusion. It has a total of five parameters: 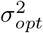 the variance of the optical noise, *n* the number of vesicles, *p* the probability of each vesicle being released, λ the scale and *γ* the shape parameters of the gamma distribution.

### G. Inferring release parameters from quantal peaks

An intuitive approach to discriminate release events from failures lies in classifying a trial as a failure of release if the observed peak fluorescence is less than twice the standard deviation of the optical noise, and success otherwise (Fig. II FA). From the distribution of success amplitudes, one then extracts the mean and coefficient of variation, called potency and CV_suc_, respectively. It is not immediately clear, however, how false positives and false negatives arising from a thresholded detection method influence the estimates of potency and CV_suc_. In this section, we use computer simulations to determine the bias introduced by optical noise on these measures.

**FIG. 4.**
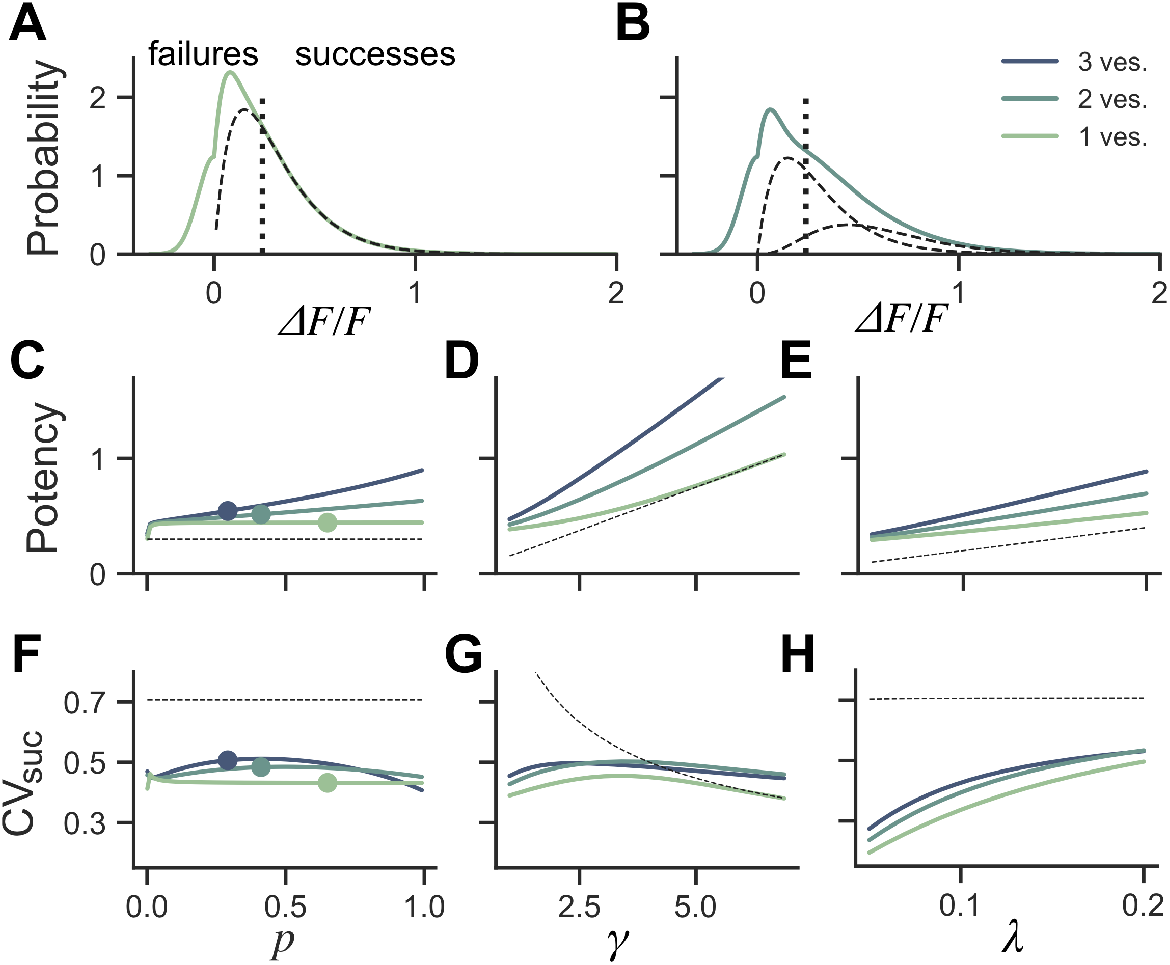
Dependence of success distribution on synaptic release properties. **A** Peak amplitude probability in the univesicular release model. All peak amplitudes occurring below the detection threshold (vertical dotted line) are classified as failures. The underlying distribution of successes (dashed black curve) shows a small portion of false negatives. **B** Peak amplitude probability in the multivesicular release model with n =2 vesicles. The distribution underlying one- and two-vesicle released are shown as dashed black curves. In A-B the probability distributions are drawn as histograms with bin size of 0.01. The mean amplitude of successes (potency) is shown as a function of the **C** the release probability for fixed shape (*γ* = 2) and scale (λ = 0.15) **D** as a function of the shape paramater *γ* for fixed release probability (*p* = 0.65) and scale (λ = 0.15) and **E** as a function of the scale parameter λ for fixed release probability (*p* = 0.65) and shape (*γ* = 2). The CV of successes is shown as a function of **F** the probability of release, **G** the shape parameter, and **H** the scale parameter. In C-H, three curves are shown for *n* = 1, 2, 3 vesicles. The dashed curve (black) shows the potency under the univesicular model in the absence of optical noise and with a detection threshold at zero.

To quantify the bias arising from classification errors, we generated surrogate amplitudes and calculated the potency and CV*_suc_* using a threshold corresponding to two standard deviations of the optical noise (Fig. II FA,B). We compared these estimates to potency and CV*_suc_* calculated without classification errors. We found that for *γ* = 2 and λ = 0.15, classification errors leads to an over-estimation of the potency for all release probabilities (Fig. II FC). This overestimate was restricted to the lower range of shape-parameter (Fig. II FD) and scale parameter values(Fig. IIFE). These biases are overall relatively small, but the effects of optical noise are more dramatic on the calculation of CV*_suc_*. Given *γ* = 2 and λ = 0.15, we found that CV*_suc_* is drastically underestimated for all *p* (Fig. II FF). This underestimate arises in a range of shape and scale parameters in the low range (Fig. II FG-H). In the case of threshold classification of successes and failures, we conclude that CV*_suc_* will be heavily underestimated when the skewness is noticeable and the quantal size (*γ*λ) is small.

Next we investigated the consequence of skewed distribution on the identification of quantal parameters *n* and *p*. Common approaches to estimate quantal parameters are based on the identification of quantal peaks [21, 23, 24, 29]. These approaches assume that the observation of a peak in the release-amplitude histogram can be read off as a quantal mode, an assumption that is often difficult to justify [14, 34, 51]. Peak identification can be even more problematic when the release components show an important skew. Indeed, we noted that mixtures of skewed distributions rarely show quantal peaks (Fig. 5). For instance, a gamma-Gaussian mixture with *n* = 2 will transition from the absence of quantal peaks (Fig. 5A) to the presence of quantal peaks (Fig. 5B) only if the skewness of the components is reduced beyond the range predicted from biophysics (Fig. 5C). These observations extend the limitations previously raised [14, 34, 51] and shows that analysis of quantal peaks is problematic especially when the distribution of univesicular release is skewed or only for a very narrow range of release probability. Since we expect a significant skew from known vesicle diameters (Fig. 3), we sought a different method for extracting release properties.

**FIG. 5.**
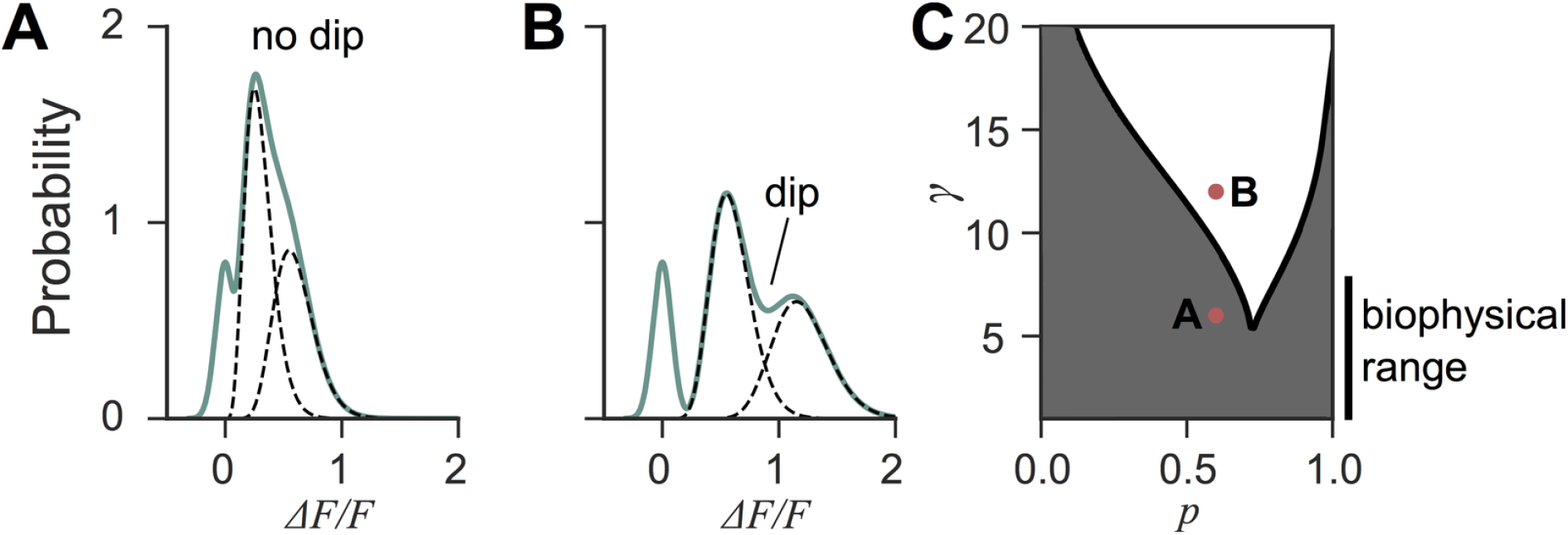
Quantal peaks are rarely apparent in mixtures of skewed distributions. **A** Peak amplitude probability density function (green curve) of the gamma-mixture with *n* = 2 vesicles, a biophysical skewness *γ* = 6 and release probability *p* = 0.6. No dip is apparent between the individual components (dashed curves). **B** Peak amplitude distribution (green curve) of the gamma-mixture model with reduced skewness, *γ* = 10. A dip (indicated) can be observed between the quantal peaks (dashed curve). **C** Phase portrait of the presence, or absence, of a dip for a model with two vesicles (*n* = 2). The presence of a dip is restricted to small skewness (*i.e*. large *γ*) and a narrow range of release probability (white region). The shaded region represents parameter value combinations not associated with a dip in the probability density function. The parameter values used in A and B are indicated with red dots.

### H. Inferring release properties using likelihood maximization

Maximizing the likelihood function provides an appealing alternative to feature-based approaches [6, 43] or quantal-peaks approaches [24]. This approach does not rely on a trial by trial classification of successes and failures. Instead, the task is to find the set of parameters (*n*, *p*, *γ*, λ, *σ*_opt_) that maximize the probability of observing all recorded amplitudes given our gamma-Gaussian model. In the case of the likelihood written in Eq. 2, there is no guarantee that only one such maximum exists, which means that it can be difficult to find the global maximum. Likelihood maximization algorithms can greatly help in solving this type of tasks.

For our problem, the Expectation-Maximization (EM) algorithm appears a natural choice since it was developed to improve parameter inference in mixture models [16]. The EM algorithm has been used previously to in fer synaptic properties, but using different experimental and computational methodologies [2]. For efficient use of this algorithm, it is critical to derive estimation formulas specific to a given problem. Since we are not aware of any such estimation formulas for the gamma-Gaussian mixture (Eq. 2), we next describe our adaptation of the EM algorithm to this context.

The likelihood maximization in the EM algorithm is associated with the principle of gradient ascent [54]. Accordingly, it begins with an initial guess, and then iteratively updates these estimates to gradually maximize the likelihood 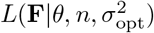 of observing the *N* observations of fluorescence amplitude denoted by the vector **F** given the parameters *θ* = (*p*, *γ*, λ). Given an initial guess *θ*_0_ = (*p*_0_, *γ*_0_, λ_0_), the algorithm will find the optimal value of each parameter 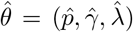. The parameter *n* will be treated as a meta-parameter to the EM algorithm, whose optimum is obtained by finding the 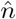 with its own optimal 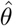 that maximizes the likelihood 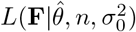, or equivalently, minimize the negative log-likelihood. The variance of the optical noise, 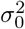, can be estimated independently by calculating the variance of the null distribution (see Methods).

Typically, a good initial estimate of the parameters can greatly speed up the inference process. In the present case, we have argued that a good prior on the shape parameter can be obtained from the biophysics of vesicle release with known, Gaussian distributed, vesicle diameters [38]. We initialize the shape parameter to a value of *γ*_0_ = 4. To initialize the probability of release, we observe that only optical noise can capture fluorescence amplitudes smaller than zero. Therefore, we compute the fraction, *c*, of the total number of observations falling below zero and equate this to half the failure probability.

This suggests the initialization *p*_0_ = 1 − (2*c*)^1/*n*^. There remains the initial value of the scale parameter. Given that the mean of fluorescence amplitude of the model is *npγ*λ, we use the mean of the observed fluorescence amplitudes *μ_F_* to initialize λ_0_ = *μ_F_*/*np*_0_*γ*_0_.

The EM algorithm is iterative and variational. That is, it first approximates the likelihood by an auxiliary function, which we will call *Q*. It then iterates between a maximization of this auxiliary function (the maximization step) and an improvement to the approximation by generating a new auxiliary function (the expectation step). Using *b*(*k|N,p*) to denote the kth binomial coefficient, the likelihood over *N* observations

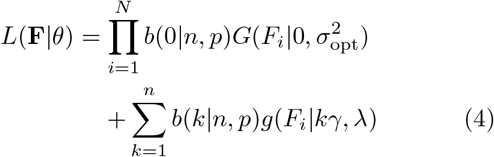

is replaced by

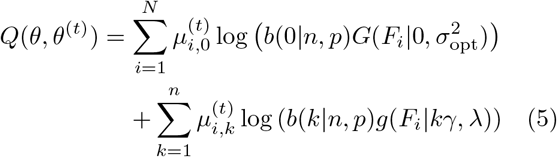

This auxiliary function relies on *N*(*n* + 1) variables 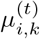. These are the posterior probabilities of sampling from the kth component given a guess of the parameters *θ*^(*t*)^.

In the expectation step, we compute the posterior

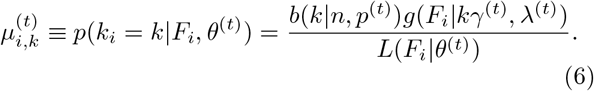

The posterior probabilities for 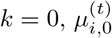, would need to be considered only if we were to use the EM algorithm to determine *σ*_opt_. Importantly, these are computed using the previous guess *θ*^(*t*)^ = (*γ*^(*t*)^, λ^(*t*)^, *p*^(*t*)^). Initially, the calculation is performed using an initial guess of the parameters, *θ*_0_.

In the maximization step, we compute the new parameter set, which maximizes the auxiliary function *θ*^(*t*+1)^ = argmax*_θ_Q*(*θ*, *θ*^(*t*)^). This is done via three re-evaluation formulas, obtained by setting the gradient of Q to zero. In what follows, we will use 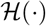 to denote the Heaviside function. The first formula gives an update of *p*

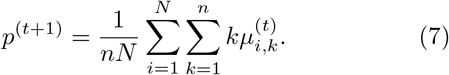

To compute the second, we first calculate the model mean

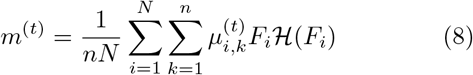

and then maximize the terms of Q that depend on *γ*^(*t*+1)^

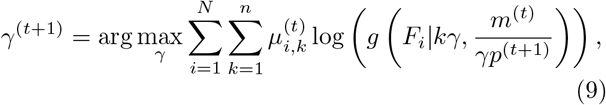

The third formula updates the scale factor

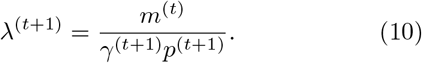

The expectation and maximization step are then repeated in alternation until convergence, which is defined by a tolerance value on the likelihood update.

We use these parameter estimates to compute the log-likelihood using Eq. 4. Repeating the EM-method for *n* within a physiological range of 1-10 allows us to find the number of vesicles 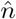 which maximizes the log-likelihood

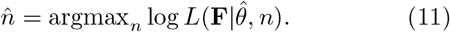

Since the results may depend on the initialization point, we repeat the procedure with ten different initialization points. The parameter values associated with the highest likelihood become our parameter estimates.

### I. Statistical inference on surrogate data

To determine the precision and the validity of the EM method for extracting release properties, we apply the method on simulated data. We assume that the fluorescence amplitude are sampled from the gamma-Gaussian distribution. Once a sample is drawn, we will use the EM method to extract the release properties, namely the parameters *γ*, λ, *N* and *p*. Knowing the true parameters allows us to calculate the average difference between estimated and true parameters (bias) and the size of random fluctuations in the estimated parameters (variance). Since these estimator bias and variance will depend on the specific set of parameter values used to generate surrogate data, we must explore different types of parameter values. For the sake of illustration, we consider three cases: i) Univesicular release (Fig. 6A), ii) multivesicular release with a low value of the shape parameter corresponding to the absence of dip in the probability distribution (Fig. 6B) and; iii) multivesicular release with a high value of the shape parameter leading to well resolved quantal peaks but inconsistent with the biophysical constraints (Fig. 6C).

**FIG. 6.**
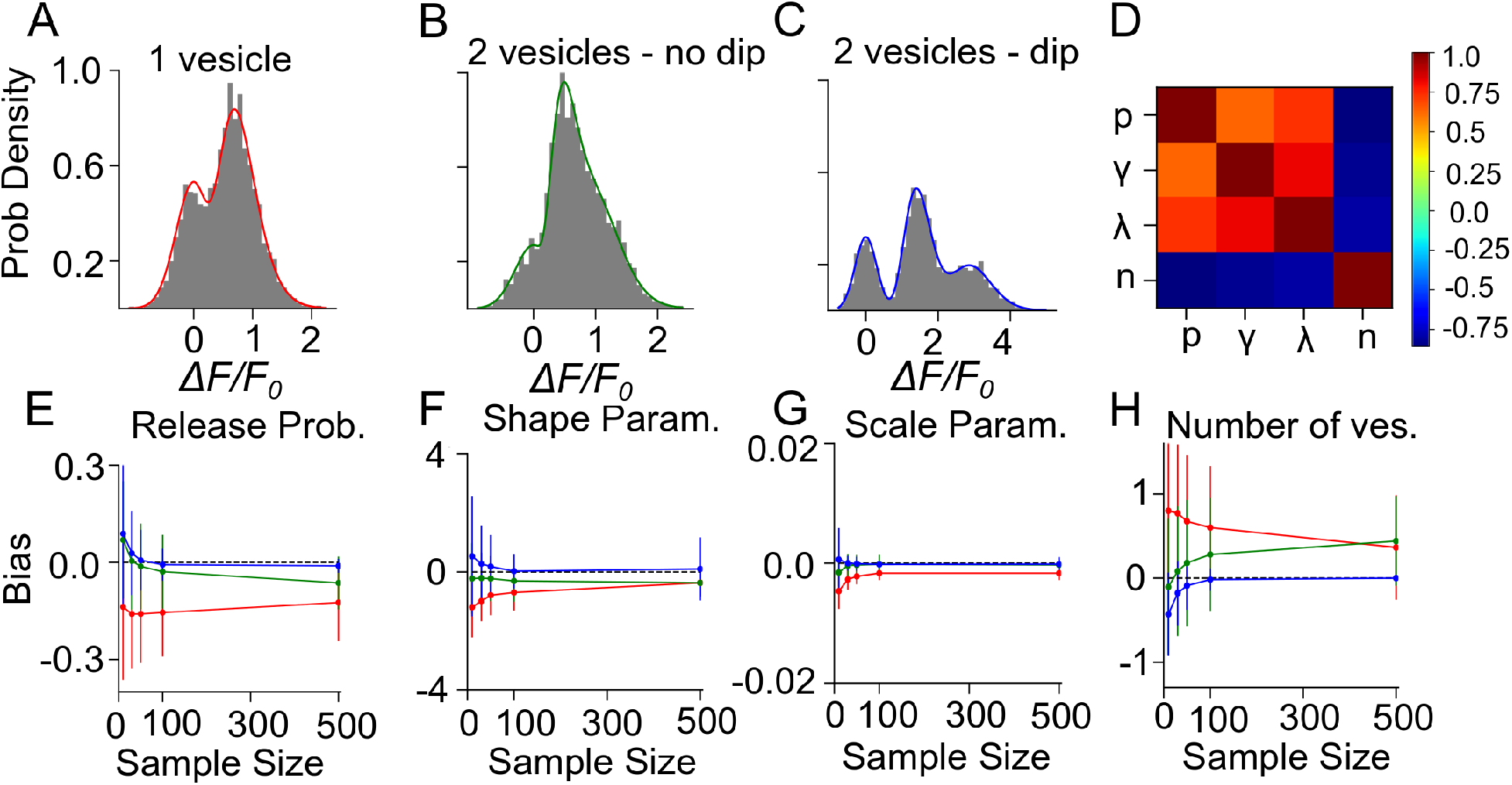
Validation of the Expectation-Maximization method on simulated gamma-mixtures. Count histograms for simulated data (gray bars) and best fit probability density function (full line) for a gamma mixture with **A** *n* = 1 vesicles, a skew *γ* = 7, scale λ = 0.12 and release probability *p* = 0.6, **B** *n* = 2 vesicles, skew *γ* = 6, scale λ = 0.1 and release probability *p* = 0.55, **C** *n* = 2 vesicles, skew γ=15, scale λ= 0.1 and release probability *p* = 0.51. **D** Correlation coefficient between parameter estimates of simulated data B with sample size = 100. The bias of estimates for **E** release probability (p), **F** shape parameter (*γ*), **G** scale parameter (λ), and **H** number of vesicles (*n*) is shown as a function of number of samples. Error bars show parameter estimates s.d.

We computed the bias and standard deviation of the estimates using 500 surrogate experiments and the expectation-maximization algorithm of the gamma-Gaussian mixture (see Methods Sect. IV E). Since both the bias and the standard deviation are expected to depend on the number of samples per dataset, *N*, we report the bias and standard deviation as a function sample size. The correlation coefficients shown in Figure 6D reveal two interactions. Firstly, release probability as well as the shape and scale parameter estimates are strongly correlated. Secondly, these three parameter estimates are anti-correlated with the estimate of the number of vesicles. These compensations are also reflected in the sample-size dependent biases, where an underestimate (overestimate) in *n* is accompanied by an overestimate (underestimate) in the other parameters, (Fig. 6E-H). This reflects the fact that *n* is determined in a separate step from the other parameters and that for a larger *n* the other parameters must decrease to keep the same mean amplitude.

Next, we consider the bias and variance of estimators for a sample sizes of 50, which represents a realistic sample size for our experimental conditions. At those sample sizes, we find that our method underestimates release probability, shape parameter and scale parameter (Fig. 6E-G, red lines). These biases reflect the fact that the number of vesicles can only be overestimated (Fig. 6H). Still considering sample size of 50 but for surrogate data with two vesicles, we see that the bias in the number of vesicles is much reduced (<0.25), and so is the bias in the release probability and the scale parameter. There remains a small underestimation of the shape parameter (Fig. 6E-G, green lines). These biases are much reduced and become negligible in the less realistic situation where quantal peaks can be identified (Fig. 6E-H, blue lines). Lastly, we note that the estimator standard deviations at *N* = 50 are sufficiently substantial to require averaging over multiple synapses in order to make precise parameter estimates.

### J. Statistical inference on experimental data

We next apply this EM algorithm on experimental data from iGluSNFR-mediated optical recordings of glutamate release. We used recordings of iGluSNFR-mediated signal induced by trains of ten axonal electrical stimulation at low frequencies (1,2,4 and 8Hz), from which we extracted a distribution of fitted release amplitudes (see Methods). Amplitude distribution from an exemplar spine is shown in Figure 7A. This distribution is captured very well by the gamma-Gaussian mixture model (Eq. 2). The best fit for this recording was achieved for shape parameter *γ* = 1.4, scale parameter λ = 0.2, release probability *p* = 0.42 and 2 vesicles. Figure 7A shows that the theoretical distribution fits the empirical histogram well. This fit arises from individual components having an important skew. The inset of Figure 7A shows the negative log-likelihood as a function of the number of vesicles. Although there is a clear minimum at *n* = 2 vesicles, the curvature is fairly large, as is predicted by the small estimator variance (Fig. 6H) under Cramer-Rao inequality. Importantly, the likelihood is considerably worse for the *n* = 1 model compared to any *n* > 1 models. Altogether, parameter inference using this EM algorithm on iGluSNFR-based analysis of glutamate release at single synapse argues that an action potential stochastically triggers the fusion of a few vesicles releasing a variable and highly skewed amount of glutamate.

**FIG. 7.**
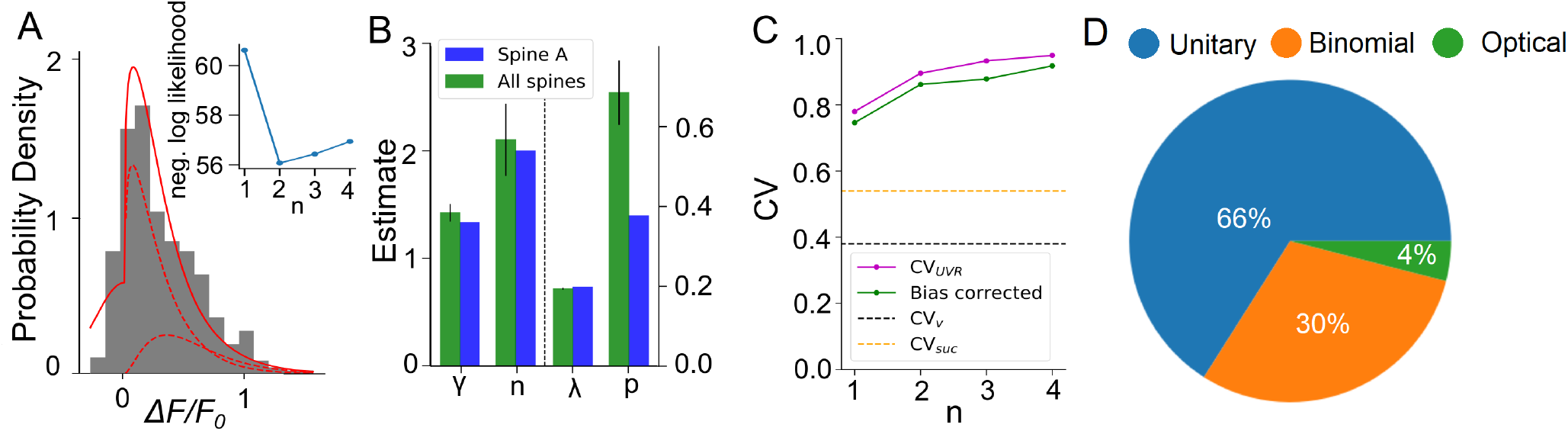
Inferring quantal parameters from IgluSNFR recordings. A Evoked fluorescence amplitude histogram for one exemplar spine (gray bars) and probability distribution of the gamma-Gaussian model with properties inferred using the EM algorithm (full red line). Individual release components for *k* = 1 and *k* = 2 are also shown (dashed red lines). Inset shows the negative log likelihood calculated by the EM algorithm versus number of vesicles released, n, for the spine shown. **B** Mean release properties obtained from the spine shown in A and a set of 18 spines. Bars left of dashed line use left axis scale, bars right of dashed line use right axis scale. Error bars represent s.e.m. The averages are 0.194 ±0.003 for λ, 0.69 ± 0.08 for p, 2.1 ± 0.3 for n and 1.42 ± 0.08 for *γ* (mean ± SEM). Error bars show SEM. **C** Univesicular CV when n is the chosen vesicular release by the EM algorithm (blue), and averaging over all estimates at that n (green). The black dashed line shows the theoretical univesicular CV. **D** Factors explaining the variance in synaptic transmission. Based on average parameters obtained in Fig. 7 and Eq. 3, we can parse out the variability in terms of optical noise (optical; green), the stochastic release of 0, 1 or 2 vesicles (binomial; orange) and the unequal potency of each vesicle (UVR; blue).

We repeated this analysis on a set of experiments from 18 different spines (Figure 7B). Here, the average number of vesicles fitted by the algorithm was 2.10 ± 0.3, while individual spines were best fit by n ranging between 1 and 3 vesicles. The average shape parameter value was 1.42±0.08. This parameter regulates the univesicular releases, the univesicular *CV_UVR_*, described in the biophysical predictions. In principle, the average shape parameter fitted by the EM algorithm should correspond to a univesicular *CV_UVR_* of 0.84, but we recall that our estimates of the shape parameter were shown to bear a small-sample bias, which we estimated to negative 0.2 (Fig. 6F). As a consequence, our bias-corrected estimate of univesicular CV is 0.77. In comparison, we predicted that a *CV_v_* = 0.38 (Fig. 3) would arise from known vesicle dimensions, thus a difference of 0.39. To see how our estimate of univesicular CV depends on the number of vesicles in the model, we fixed *n* and inferred *CV_UVR_* for each spine (Fig. 7C). We find that increasing *n* increases the *CV_UVR_* inferred. This CV*_UVR_* remained high and above both the variability expected from volumes (CV*_v_*) and the variability of thresholded successes (CV*_suc_*). This is consistent with the view that CV*_v_* is a lower bound and CV*_suc_* is underestimated (Fig. II F). In sum, statistical inference of our gamma-Gaussian model suggests CA3-CA1 synapses stochastically release 1-3 vesicles with variable quantum.

The formalism outlined above allows to begin parsing out the variability of synaptic transmission at single synapses. Using the average parameters extracted using the expectation-maximization algorithm, Eq. 3 can be used to separate the variability of observed evoked amplitudes in three terms. The first term captures the variability due to the glutamate sensor itself and to concurrent optical measurements The second term captures the fluctuations in release amplitude attributable to a single vesicle release and scaled by the average number of vesicle released (i.e. diffusion, loading and vesicle volume). The last term captures the variability of releasing sometimes two, one or zero vesicles (with zero unitary variability). We named these sources of variability optical, unitary and binomial, respectively. As shown in Figure 7D, we estimate that that 4% of the variability was optical, 30% binomial and 66% was unitary. Thus the results suggests that, despite the fluctuating number of vesicles being released, the variability of synaptic transmission arises mostly from the variability in unitary vesicle content released.

## III. DISCUSSION

The use of the glutamate fluorescent reporter iGluSnFR provides a valuable proxy of glutamate release at single central synapses. To interpret the variability of glutamate release observed in our recordings, we built a gamma-Gaussian mixture model based on stochastic release of vesicles, each with a variable diameter size and additional sources of variability. We highlighted important biases measures of the variability of successes based from threshold classification methods and provided an alternative method based on the expectation-maximization algorithm. Surrogate data analysis revealed that our statistical method has relatively small biases and allows inference of estimates of quantal parameters. Together, these experimental and analytical tools allow to resolve the dynamic structure of synaptic transmission.

Several experimental methodologies have been developed in order to monitor transmission at single synapses. One main advantage of optical methods over classical electrophysiological techniques is that the experiment is localized to an unambiguous source spine and, presumably, synaptic contact. While strong criteria have been developed in the past to classify electrophysiological experiments that are likely to arise from a single synaptic contact (i.e., minimal stimulation criteria) [17, 18, 30, 39, 47], some of these criteria may introduce selection bias in population sampling, favouring against synapses that display multi-quantal and/or a high probability of release. Another distinct advantage of using iGluSNFR, in particular, as a postsynaptic reporter of synaptic release is the non-reliance on post-synaptic glutamate receptor activation. iGluSnFR can report glutamate release events in the presence of glutamate receptor antagonists (Figure 2I), which offers experimental flexibility. Moreover, interpretation of quantal events recorded using either calcium-based optical methods or electrophysiological methods are confounded by issues such as the non-linear relationship between glutamate concentration and glutamate receptor conductance [44], the non-uniform distribution of glutamate receptors in the postsynaptic membrane[7], and the distance between the release site location and the postsynaptic nanodomain of glutamate receptor density [19]. iGluSnFR, being plasma membrane localized without anchoring domains, is presumably evenly distributed in the postsynaptic membrane and present in close proximity to the site of glutamate release. Also, as demonstrated in Figure 3F, iGluSnFR provides a linear report of physiologically-relevant glutamate release. Although only superficially dealt with here, the faster kinetics of iGluSNFR transients widens the ability to study dynamic regulation of release during trains of stimulation, when compared to the convolved proxy of glutamate release based on NMDAR-dependent calcium detection. Altogether, this method addresses a number of historical limitations and provides a welcome complement to existing methodologies to study basic features of synaptic transmission and plasticity.

Synaptic transmission is variable. Obtaining an accurate estimate of the size of this variability is an obligatory step in order to parse information content from noise during neural communication. Applying a traditional threshold-based classification of successes and failures on iGluSnFR transientsm we obtained a fairly low average variability CV*_suc_* ≈0.5 (Fig. 7C). Some of our recordings showed CV_suc_ in the 0.2-0.4 range, which closely matches the values reported for putative single-synapse electrophysiological recordings using either manual or threshold-based classification of failures and successes [5, 17, 20]. We however readily observed synapses that showed a higher CV*_suc_* (up to 0.8) when optically probed. It is likely that these synapses would have been ignored when applying selection criteria commonly used for minimal stimulation experiments. Such threshold-based classification, however, inherently introduces classification errors, which can dramatically alter estimates of CV*_suc_*. The statistical methodology presented here should circumvent this issue. Consistent with our estimates on surrogate data (Fig. II F), we observed that the variability of individual synaptic release can be much higher CV*_UVR_*=0.8. Further consistent with the effect of classification errors, our estimate of the average release probability of individual release is higher using the expectation algorithm (p = 0.69; Fig. 7B) than using a threshold-based approaches (previous estimates were <0.61 [20], 0.4 [5] and 0.2-0.4 [17]). While further studies are required to further validate these estimates, our results suggest that individual synaptic release often yields very small but non-zero release and a variability considerably higher than previously thought.

## IV. METHODS

The essential elements of optical quantal analysis are described in the main text. In this section, we give additional precisions on both experimental and computational methods.

### A. Organotypic Slices and Biolistic Transfection

A detailed description of our methodology for hippocamal organotypic slice preparation and biolistic transfection is described in Soares et al. (2014) [46]. Briefly, organotypic slices were prepared from Sprague Daley rats (Charles River Laboratories, MA, USA) using the interface method originally described in Stoppini *et al*. 1991 [48]. Animals were anesthetized in an isofluorane infused chamber, decapitated, and hippocampi were removed in ice cold cutting solution containing (in mM): 119 choline chloride, 2.5 KCl, 4.3 MgSO_4_, 1.0 CaCl_2_, 1.0 NaH_2_PO_4_-H_2_O, 1.3 Na-ascorbate, 11 glucose, 1 kynurenic acid, 26.2 NaHCO_3_, saturated with 95% O_2_ and 5% CO_2_ (pH = 7.3; 295-310 mOsm/L). Transverse slices were cut at 400 *μ*m thickness using a MX-TS tissue slicer (Siskiyou, Grants Pass, OR) and cultured on 0.4 mm millicell culture inserts (EMD Millipore, Etobicoke, Canada) at a temperature of 37°C. Hippocampal slices were transfected at 6-7 DIV using a hand held gene gun (Biorad, Hercules, CA). Cartridges for the gene gun were prepared in advance by precipitating 50 *μ*g of cDNA plasmid onto 8-10 mg of gold microparticles (1.0 *μ*m diameter; Biorad) at a ratio of 80/20 by weight of either iGluSNFR or GCaMP6f and mCherry cDNA plasmid, respectively. The precipitation step was performed in a 0.1 M KH_2_PO_4_ buffer solution containing 0.05 mM protamine sulfate (rather than spermine, as per previous protocols). The DNA-gold precipitate was washed and suspended (3 times) in 100% ethanol before loading in the tubing station. Once the cartridges were dried and cut, they were placed in a sealed container with desiccant pellets at 4 °C until used. The DNA-coated gold particles were delivered to the slice using ~180 psi of helium air pressure. A modified gene gun barrel was used to protect slices from helium blast [46]. Imaging experiments were performed 3-5 days after biolistic transfection.

### B. Optical Recording of IGluSNFR transients

Slices were removed from culture and placed in a custom recording chamber under a BX61WU upright microscope (60X, 1.0 NA objective; Olympus, Melville, NY). Slices were continuously perfused with a Ringer’s solution containing (in mM): 119 NaCl, 2.5 KCl, 4 MgSO_4_- 7H_2_0, 4 CaCl_2_, 1.0 NaH_2_PO_4_, 11 glucose, 26.2 NaHCO_3_ and 1 Na-Ascorbate, saturated with 95% O2 and 5% CO_2_ (295-310 mOsm/L). For evoked stimulation experiments, a glass monopolar electrode filled with Ringer’s solution was positioned adjacent to transfected cells in the direction of CA3. Simultaneous two-photon imaging of iGluSnFR and mCherry was performed using a Ti:Sapphire pulsed laser (MaiTai-DeepSee; Spectra Physics, Santa Clara, CA) tuned to 950 nm. Emission photons were spectrally separated using a dichroic mirror (570 nm) and the emitted light was additionally filtered using two separate bandpass filters (iGluSNFR: 495-540; mCherry: 575-630). The sampling frequency of our line-scan experiments depended on length of the imaged line segment (drawn over a spine and its parent dendrite), but was typically in the range of 1.2 - 1.5 ms / line for all optical quantal analysis experiments. This sampling rate was more than sufficient to fully capture and quantify the rise and falling phases of iGluSnFR transients. In our hands, an optimal trade-off between signal-to-noise, sampling frequency, and reduced bleaching, was obtained by using a 4 *μ*s pixel dwell time. In the frame scan configuration, the sampling limit of our optical system was 65 ms/frame (2 *μ*s pixel dwell time; 256 × 256 pixel frame) when scanning bidirectionally, which was sub-optimal for optical quantal analysis.

Surveying methodology was designed to increase the probability of finding dendritic spines that were responsive to the electrical stimulus. While our frame scan configuration offered the spatial resolution to monitor several spines at once, we found it difficult in practice to identify rapid iGluSnFR-mediated transients due to a low signal to noise ratio an infrequent sampling rate. As a result, line scans were exclusively used to survey the dendritic arbor for responsive spines. Short duration (0.1 ms) low intensity (5-25 mA) stimuli were delivered to the slice at low-frequency (0.1 Hz) while randomly surveying dendritic spines in the apical arbor of transfected cells. To facilitate the process of finding a responsive spine, line scans were performed simultaneously through multiple nearby dendritic spines and, generally, a paired-pulse stimulus (50-100 ms inter stimulus interval) was delivered to increase the probability of detecting glutamate release. Dendritic spines that were unresponsive to an initial probing phase consisting of 5-10 paired pulse stimuli, were not considered for further analysis, while spines demonstrating responsiveness to these initial probing stimuli were selected for quantal analysis experiments. Fluorescent transients were resolvable by eye and on-line analysis was not necessary. Prior to starting an optical quantal analysis experiment at a responsive spine, the stimulus intensity was gradually reduced up to a minimum where time-locked responsiveness was still observed. The process of identifying a responsive spine was generally not trivial and often necessitated several re-positioning of the stimulating electrode. Once a responsive spine was found, however, it was extremely rare to lose fluorescent responsiveness in response to electrical stimulation during an experiment.

### C. Whole-cell electrophysiology and two-photon glutamate uncaging

Whole-cell recordings were carried out using an Axon Multiclamp 700B amplifier. Electrical signals were sampled at 10 kHz, filtered at 2 kHz, and digitized using an Axon Digidata 1440A digitizer (Molecular Devices, Sunnyvale, CA). Transfected CA1 pyramidal neurons were targeted and patched using borosilicate glass recording electrodes (World Precision Instruments, Sarasota, FL) with resistances ranging from 3-5 MΩ. All uncaging evoked currents were recorded at a holding potential of −70 mV. The intracellular recording solution contained (in mM): 115 cesium methane-sulfonate, 0.4 EGTA, 5 tetraethylammonium-chloride, 6.67 NaCl, 20 HEPES, 4 ATP-Mg, 0.5 GTP, 10 Na-phosphocreatine (all purchased from Life Technologies, Carlsbad, CA) and 5 QX-314 purchased from Abcam (pH = 7.2-7.3; 280-290 mOsm/L). The extracellular solution was similar to the one described above but also contained 2. mM MNI-glutamate-trifluoroacetate (Femtonics, Budapest, Hungary) and a reduced concentration of MgSO_4_-7H_2_0 (1.3 mM) and CaCl_2_ (2.5 mM). For glutamate uncaging experiments, a second laser line tuned to 720 nm was used to deliver 1 ms light pulses to the tips of dendritic spines while the other laser was tuned to 950 nm to image the uncaging-evoked iGluSnFR transients.

### D. Regression for amplitude extraction

We describe the use of a template to extract the amplitude of the evoked responses. The method involves two steps. First we extract a template time-course k by computing the trial-averaged fluorescence response that is triggered by the electrical stimulation. This template is discretized, starts at the stimulation time and ends at a pre-defined time *T* after it. Trial averaging is performed on responses sufficiently isolated in time to be exempt from other synaptic events. For each trial *i* the template is scaled by *β* chosen so as to minimize the mean-squared error with the observed fluorescence 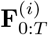 in the corresponding time window indicated by the subscript 0 : *T*. The solution of this least-square problem is well known and follows

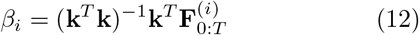

In order to report the maximum of the evoked waveform, we scale *β_i_* by the maximum value of the template. This is the value reported in Fig. 5. We calculated the null distribution by fitting the template on fluorescence measurements without electric stimulation. The variance of the null distribution serves as our estimate of measurement noise 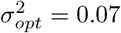.

### E. Surrogate data analysis

To generate surrogate data, we simulate *n* surrogate fluorescence amplitudes and estimate the parameter values. Each surrogate experiment is repeated *M* times in order to have *M* sets of parameter estimates 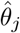. Using this set of surrogate experiments, we can compute the bias, the variance and the correlation coefficients of the estimates. The bias is calculated by averaging the difference between the estimated parameter and the simulated parameters

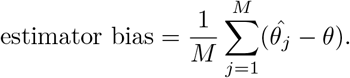

In this way, a bias greater than zero means that the parameter tends to be overestimated, and a bias smaller than zero means that the inferred parameters are erroneously small.

To estimate the precision of the estimates, we calculate the variance.

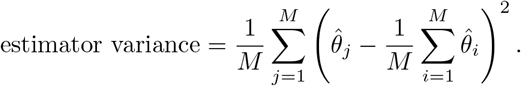

A small variance means that the estimate is precise, although it may or may not be valid.

In addition, we compute the correlation coefficient between different parameter estimates. The correlation coefficient between a parameter *θ*^(*r*)^ and *θ*^(*q*)^ is simply

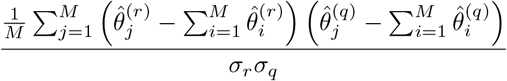

where *σ_r_* and *σ_q_* are the square root of the estimator variances for *θ*^(*r*)^ and *θ*^(*q*)^, respectively. When the correlation coefficient is positive, it means that estimation errors tend to covary with the same sign, and when the correlation coefficient is negative, it means that an estimation error in parameter *r* is associated with an estimation error of opposite sign in parameter *q*.

## V. ADDITIONAL REQUIREMENTS

For additional requirements for specific article types and further information please refer to Author Guidelines.

## AUTHOR CONTRIBUTIONS

CS, JCB, AL and RN conceived the study. CS performed the experiments. CS, DT and RN analyzed the data and performed the simulations.

